# Regulation of the error-prone DNA polymerase polκ by oncogenic signaling and its contribution to drug resistance

**DOI:** 10.1101/316729

**Authors:** Kelsey Temprine, Erin M Langdon, Krisha Mehta, Averill Clapp, Richard M White

## Abstract

Mutations in the proofreading domains of the replicative DNA polymerases polδ and polε are associated with elevated mutation rates in cancer, but the roles of other DNA polymerases in tumorigenesis remain poorly understood. One such polymerase is polκ, an enzyme that plays a key role in translesion synthesis. polκ contributes to cell survival in the face of DNA damage but can be highly mutagenic due to lack of a proofreading domain. Here we demonstrate that cancer cells under stress from oncogene inhibition upregulate polκ and shift its localization from the cytoplasm to the nucleus. This effect can be phenocopied by mTOR inhibition or glucose deprivation, analogous to stress-induced mutagenesis in *E. coli* whereby cell stress and nutrient deprivation can upregulate and activate DinB/pol IV (the bacterial orthologue of polκ). We find that cancer cells normally sequester polκ in the cytoplasm via exportin-1, likely to prevent excess mutagenesis from the error-prone nature of this polymerase. Subverting the normal nuclear-cytoplasmic shuttling by forced overexpression of nuclear polκ increases resistance of melanoma cells to the BRAF^V600E^ inhibitor vemurafenib. This data suggests a mechanism by which cancer cells regulate the expression and localization of the error-prone polymerase polκ, abrogation of which can contribute to drug resistance.

**One Sentence Summary:** Cancer cells under stress from oncogene or mTOR inhibition dysregulate the error-prone DNA polymerase polκ, which contributes to drug resistance in melanoma cells.

## Introduction

Errors in DNA replication can lead to increased mutation rates, thereby contributing to cancer pathogenesis. For example, somatic or germline mutations in the proofreading domain of DNA polymerase delta (polδ) or epsilon (polε) can lead to tumors with markedly increased numbers of point mutations (*1–3*). Aside from these two main replicative polymerases, a number of other DNA polymerases have been identified that may contribute to cancer initiation or progression (*4*). For example, inactivation of DNA polymerase eta (polη) is associated with xeroderma pigmentosum variant (XP-V), which predisposes patients to UV-induced skin cancers (*5*). Additionally, DNA polymerase iota (polι) is upregulated in esophageal squamous cell cancer, and its expression levels positively correlate with lymph node metastasis/clinical stage (*6*).

The roles of other DNA polymerases in this process are less well understood but likely could contribute to tumor progression. One such polymerase is DNA polymerase kappa (polκ), which is a member of the Y-family of DNA polymerases that plays an essential role in the DNA damage tolerance process of translesion synthesis (*7, 8*). Several previous studies have shown that overexpression of polκ can contribute to tumorigenesis and drug resistance in cancer (*9–12*). For example, overexpression of polκ in glioblastoma cells increases resistance to the DNA-damaging agent temozolomide (*12*), and it has also been found to be significantly overexpressed in lung cancer (*9*).

polκ can replicate DNA in both an error-free and error-prone manner during translesion synthesis (*13*). It can bypass thymine glycols in a relatively error-free manner (*14*), whereas it bypasses N-2-acetylaminofluorene adducts in a more error-prone manner (*15*). When replicating on undamaged DNA, polκ has a markedly high error rate due to a relatively large active site and lack of a proofreading domain (*16*). Using *in vitro* assays, it has been shown to have error rates as high as 1 error per 200 base pairs when replicating on undamaged DNA (*17*). For this reason, it is considered an “error-prone” polymerase that can induce untargeted mutations while acting either directly at the replication fork or by filling in post-replication gaps (*18*). The range of errors introduced by polκ span virtually all substitutions, although to differing degrees (with a high rate of T→G substitutions), as well as a preponderance of deletions (*16*). These error rates are substantially higher than that found for the replicative polymerases polδ and polε.

Because dysregulated polκ can be mutagenic at high levels, it is important that cells limit both its expression and access to DNA. In bacteria, this regulation is enacted via the SOS/DNA damage response along with the RpoS/starvation stress response (*19, 20*). In work spanning several decades, it has been observed that *E. coli* use a mechanism referred to as stress-induced mutagenesis to temporarily increase their mutation rate under periods of stress (*20–23*). This hypermutation is enacted as part of double-strand DNA break repair, which becomes mutagenic due in part to the activity of DinB (*24*), the *E. coli* orthologue of human polκ. This mutagenic process is regulated at three levels, as recently reviewed (*22*): 1) a double-strand break (*25, 26*), 2) activation of the SOS DNA damage response (*21*), and 3) activation of the generalized sigma S (RpoS) stress response (*20*). The SOS response, when coupled with this stress response, allows first for upregulation of DinB (*27*) and then subsequent usage of this error-prone polymerase for mutagenic repair, which results in the base substitutions and indels that are commonly observed (*20*). It is likely that deficiencies in mismatch repair contribute to this process since overexpression of MutL inhibits mutation in stationary phase but not during growth (*28*), whereas both MutS and MutH can be downregulated in part by the stress response RpoS pathway (*29*).

In contrast to the work in *E. coli*, the mechanisms regulating the expression and localization of polκ in mammalian cells remains poorly understood. In normal human tissues, polκ is widely expressed at the mRNA level (*30*), whereas in the mouse, it is highly enriched in the adrenal cortex and testis (*31*). In the mouse, protein expression using a peptide-generated antibody was noted in adrenal cortex, pachytene cells in meiosis I, post-meiotic spermatids, and some epithelial cells in the lung and stomach (*31*). At the cellular level, numerous studies using overexpression of EGFP-polκ fusion proteins have demonstrated that polκ is strongly enriched in the nucleus (*32, 33*). However, antibody staining of endogenous polκ protein using antibodies generated with either peptide fragment or full-length proteins have shown variable expression in both the cytoplasm as well as the nucleus (*31*). Analysis of the polκ promoter has shown consensus binding sites for Sp1 and CREB, both of which have been reported to transcriptionally activate its expression (*34, 35*).

Although recent observations demonstrate that polκ is mutagenic and can promote tumorigenesis and drug resistance (*9–12*), it is unknown what regulates its expression in cancer. Ectopic expression of polκ allows it to become part of the replication machinery, even in the absence of external stress, indicating that high levels of it alone may be sufficient to induce new mutations (*32*). This has important clinical implications since dysregulation of polκ expression could therefore contribute to tumorigenic phenotypes by affecting its normal subcellular localization. In this study, we demonstrate that deprivation of oncogenic signaling in melanoma, lung, and breast cancer cell lines upregulates polκ and confines it to the nucleus. These pathways converge on PI3K/mTOR signaling, a central regulator of nutrient status in the cell (*36*) that may be analogous to the generalized stress factor sigma S in *E. coli* (*22*). When cells are nutrient replete and have intact mTOR signaling, polκ is primarily present in the cytoplasm; when cells are starved or mTOR is inhibited, polκ shifts primarily to the nucleus, suggesting that the cell can dynamically regulate polκ in response to cell stress. In line with this, we find that polκ can be rapidly exported back out of the nucleus via the nuclear export machinery, implying that cells may normally use nuclear export to prevent excess mutagenesis. When this dynamic regulation is subverted by forced nuclear overexpression of polκ, we find that this increases resistance to the BRAF^V600E^ inhibitor vemurafenib in melanoma cells. Our data suggest a mechanism by which mammalian cancer cells regulate the levels and localization of the error-prone DNA polymerase polκ, dysregulation of which can contribute to drug resistance.

## Results

### MAPK inhibition induces polκ mRNA upregulation and changes the subcellular localization of its protein

Given the role of stress in upregulating DinB/pol IV in *E. coli*, we first asked whether cell stress regulated polκ expression in cancer. We reasoned that drugs blocking oncogenic drivers would induce cell cycle arrest and ultimately apoptosis and would represent an extreme form of cell stress. In human melanoma, the most common activating mutation occurs at BRAF^V600E^, which activates downstream MAP kinase signaling via MEK/ERK signaling. This provides the melanoma cells with a significant growth advantage. Small molecules targeting the BRAF/MEK/ERK pathway have been developed, some of which are being clinically used to treat melanoma patients (*37–39*). BRAF inhibitors such as vemurafenib can induce cell stress prior to overt death of the cancer cell, which can occur via the ER stress pathway (*40*) and is associated with stress-induced senescence (*41*). Based on this, we first asked whether inhibition of the MAP kinase pathway could induce polκ expression in melanoma.

To assess this, we treated the BRAF^V600E^mutant melanoma A375 cell line with the BRAF^V600E^kinase inhibitor PLX4032 (vemurafenib) (*37*) and measured polκ by qRT-PCR from 2-72 hours post exposure. This revealed an upregulation in expression of polκ that peaks at 24 hours and is sustained thereafter (Fig. 1A). We next examined expression of polκ at the protein level under similar conditions. We utilized an antibody raised against full-length human polκ protein and verified its specificity using shRNA knockdown and overexpression of endogenous polκ (Fig. S1, A to C). Although polκ has been largely reported to be nuclear localized when overexpressed as an EGFP-tagged fusion protein (*32, 33*), we surprisingly found that the Western blot using an antibody raised against the full-length protein showed that endogenous polκ was expressed in both cytoplasmic and nuclear fractions (Fig. 1B). This expression pattern is similar to that of the related family member pol eta (polη), where both cytoplasmic and nuclear expression is seen (Human Protein Atlas (*42*)). While treatment with vemurafenib did not change overall protein levels of polκ, it instead induced a specific increase in nuclear polκ and corresponding decrease in cytoplasmic polκ (Fig. 1, B and C). We confirmed this redistribution using immunofluorescence, where acute exposure to PLX4032 induced a shift of polκ from the cytoplasm to the nucleus (Fig. 1, D and E). Cytoplasmic localization of a DNA polymerase that can shift from the cytoplasm to the nucleus has been reported previously (*43–45*), suggesting that this mode of regulation might be relevant for polκ under physiologic conditions. Because BRAF activates downstream MEK/ERK signaling, we also examined whether downstream inhibitors would elicit similar effects. In two different BRAF^V600E^mutant melanoma cell lines (A375 and SK-MEL28), we found that both MEK and ERK inhibitors (*38, 39*) produced an upregulation of polκ mRNA and a nuclear shift of polκ that are similar to that seen after BRAF inhibition (Fig. 1, F to I and Fig. S2, A to D).

**Fig. 1.**
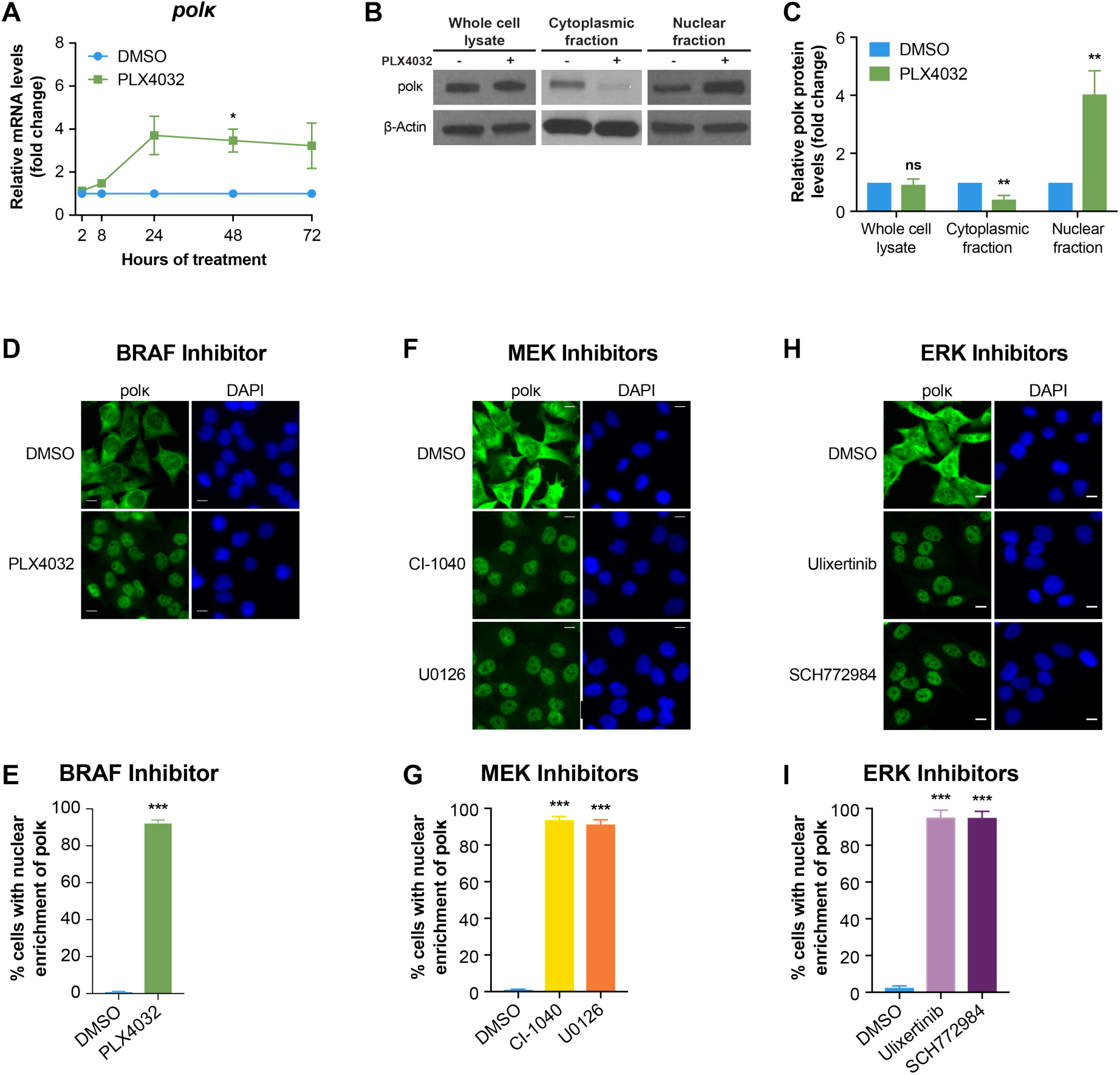
Treatment of melanoma cells with BRAF or MAP kinase inhibitors modulates polκ expression and localization. (**A**) qRT-PCR to detect the mRNA expression of polκ relative to the DMSO control was performed on A375 cells treated with DMSO or 5 µM PLX4032 for 2, 8, 24, 48, or 72 hours, Mean ± S.E.M, n=3 experiments, *P < 0.05, paired two-tailed t-test. (**B**) Western blot of A375 cells treated with DMSO (−) or 5 µM PLX4032 (+) for 48 hours. (**C**) Quantification of the Western blot data relative to the DMSO control from 5–8 independent experiments, Mean ± S.E.M, ns = non-significant, **P < 0.01, paired two-tailed t-test. β-actin or Lamin B1 served as the loading control. (**D**) Immunofluorescence staining of A375 cells treated with DMSO or 5 µM PLX4032 for 24 hours, scale bar 10 µm. (**E**) Quantification of the percentage of cells with nuclear enrichment of polκ, Mean ± S.E.M, at least 2500 cells were counted for each sample across 22 fields of view, ***P < 0.001, paired two-tailed t-test. (**F**) Immunofluorescence staining of A375 cells treated with DMSO or MEK inhibitors (10 µM CI-1040, 10 µM U0126) for 24 hours, scale bar 10 µm. (**G**) Quantification of the percentage of cells with nuclear enrichment of polκ, Mean ± S.E.M, at least 1000 cells were counted for each sample across 18 fields of view, ***P < 0.001, paired two-tailed t-test. (**H**) Immunofluorescence staining of A375 cells treated with DMSO or ERK inhibitors (1 µM Ulixertinib, 1 µM SCH772984) for 24 hours, scale bar 10 µm. (**I**) Quantification of the percentage of cells with nuclear enrichment of polκ, Mean ± S.E.M, at least 250 cells were counted for each sample across 6 fields of view, ***P < 0.001, paired two-tailed t-test.

### DNA damage or cell cycle arrest are not sufficient for polκ localization

The induction of DinB in bacteria relies upon two related systems: the SOS/DNA damage response and the RpoS-controlled general/starvation stress response. We reasoned that the nuclear localization of polκ in response to MAP kinase inhibition might act through one of these two mechanisms. To test this, we treated A375 melanoma cells with vemurafenib and then checked for markers of DNA damage (*46*), including gamma H2AX (γH2AX), 53BP1, phosphorylated Chk1 (p-Chk1), and phosphorylated Chk2 (p-Chk2) (Fig. S3A) but saw no induction of any of these markers. We also tested whether knockdown of p53, which is a regulator of the DNA damage response pathway (*47*), would prevent the observed subcellular shift but failed to see any change in the nuclear localization of polκ after treatment with PLX4032 (Fig. S3, B to D). Another possibility was that cell cycle arrest was the signal, as would be expected during a starvation response or during MAPK inhibition. To test this, we treated A375 melanoma cells with either the CDK4/6 inhibitor PD0332991 (*48*) or the BRAF^V600E^inhibitor vemurafenib and then assessed the cell cycle using flow cytometry and polκ localization using immunofluorescence. As expected, both of these interventions led to a near complete G1 arrest; however, treatment with the CDK4/6 inhibitor resulted in little to no nuclear polκ (Fig. S4, A to C). Taken together, these data suggested to us that other mechanisms might be responsible for the induction of polκ.

### mTOR inhibition rapidly induces polκ nuclear accumulation

As we saw no effect of cell cycle inhibition or DNA damage, we next turned to mTOR signaling, a central sensor and effector of growth factor signaling, nutrient status, and stress (*36*). The mTOR pathway can be activated by RAF/MEK/ERK signaling either directly or via cross-pathway activation (*49–52*). In addition, BRAF inhibition with vemurafenib has recently been shown to induce the ER stress response (*40, 53*), which is associated with suppression of mTOR signaling (*54*). Based on this, we reasoned that MAPK inhibition might be acting to dampen downstream mTOR signaling to mediate the effect on polκ. To test this, we treated A375 melanoma cells with inhibitors of BRAF/MEK/ERK signaling for 24 hours and then measured mTOR pathway activation via levels of phosphorylated S6 (p-S6), a target of S6K (*36, 55*). We found that these MAPK inhibitors potently decreased levels of p-S6 (Fig. 2, A and B and Fig. S5, A and B), and that this inversely correlated with the shift of polκ to the nucleus: cells with the lowest level of p-S6 had the highest levels of nuclear polκ (Fig. 2C). These observations prompted us to then test whether inhibitors of the mTOR pathway itself would lead to an effect on polκ. We treated A375 melanoma cells with several inhibitors of the PI3K/mTOR pathway, including mTOR inhibitors (rapamycin or PP242) or a PI3K inhibitor (LY29002). We found that these inhibitors potently induced nuclear polκ to a level comparable to that seen with the BRAF/MEK/ERK inhibitors (Fig. 2, D and E), but it occurred much more rapidly. Whereas the MAP kinase inhibitors took ~24 hours for full induction of nuclear polκ, the mTOR pathway inhibitors could do this in as little as ~6 hours.

**Fig. 2.**
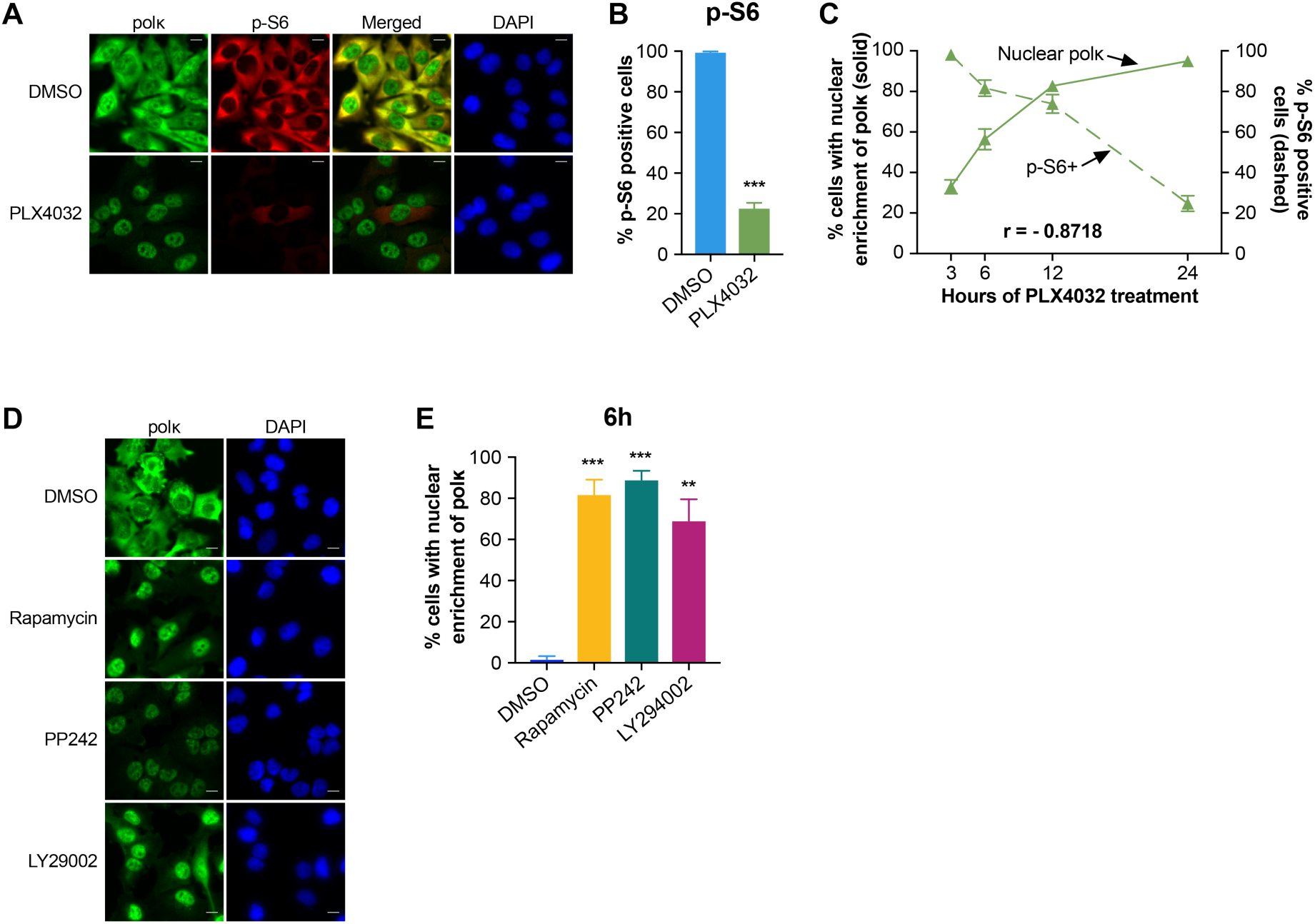
mTOR signaling regulates polκ subcellular localization. (**A**) Immunofluorescence staining of A375 cells treated with DMSO or 5 µM PLX4032 for 24h hours, scale bar 10 µm. (**B**) Quantification of the percentage of cells with phosphorylated S6 (p-S6, S240/244), Mean ± S.E.M, at least 400 cells were counted for each sample across 8 fields of view, ***P < 0.001, paired two-tailed t-test. (**C**) Immunofluorescence staining of A375 cells treated with 5 µM PLX4032 for 3, 6, 12, or 24 hours. The bar graph shows the percentage of cells with nuclear enrichment of polκ (left axis, solid line) and the percentage of cells with phosphorylated S6 (p-S6, right axis, dashed line), Mean ± S.E.M, at least 150 cells were counted for each sample across 4–8 fields of view. The Pearson correlation coefficient (r) was calculated for the percentage of cells with nuclear enrichment of polκ vs. the percentage of cells with phosphorylated S6. (**D**) Immunofluorescence staining of A375 cells treated with DMSO, 0.5 µM rapamycin, 5 µM PP242, or 30 µM LY294002 for 6 hours, scale bar 10 µm. (**E**) Quantification of the percentage of cells with nuclear enrichment of polκ after 6 hours of treatment, Mean ± S.E.M, at least 200 cells were counted for each sample across 8–12 fields of view, **P < 0.01, ***P < 0.001, paired two-tailed t-test.

### polκ dysregulation occurs in other cancer types

We next wished to determine if the observed effects on polκ were specific to melanoma/the BRAF pathway, or if they could be applied more broadly. To test this, we examined polκ mRNA levels and subcellular localization in breast and lung cancer cell lines, which harbor unique oncogenic dependencies that differ from melanoma. PC-9 lung cancer cells harbor activating mutations in EGFR and are critically dependent upon that growth factor signaling pathway. We treated PC-9 cells with either the EGFR inhibitor erlotinib (*56*) or the BRAF^V600E^inhibitor PLX4032 (which we would expect to have no effect as these cells contain wild-type BRAF) and then measured the expression of polκ and determined its localization. Both high and low doses of erlotinib caused induction of polκ mRNA and a shift of polκ to the nucleus in the PC-9 cells, but as expected, this effect was not seen with PLX4032 (Fig. 3, A and B and Fig. S6A). Analogous to what we observed for melanoma, this shift of polκ to the nucleus was accompanied by a loss of p-S6 in the lung cancer cells (Fig. 3, A and C). A similar effect was seen in a breast cancer cell line. SK-BR3 cells, which overexpress the HER2 oncogene, were treated with the HER2 inhibitor lapatinib (*57*) or the BRAF^V600E^inhibitor PLX4032. Lapatinib caused a marked induction of polκ expression, nuclear accumulation of polκ, and suppression of p-S6 (Fig. 3, D to F and Fig. S6B), an effect that was not seen with PLX4032 in this cell line as expected. These results demonstrate that the observed effects on polκ are specific to the driver oncogene, and that they can be generalizable to multiple cancer types.

**Fig. 3.**
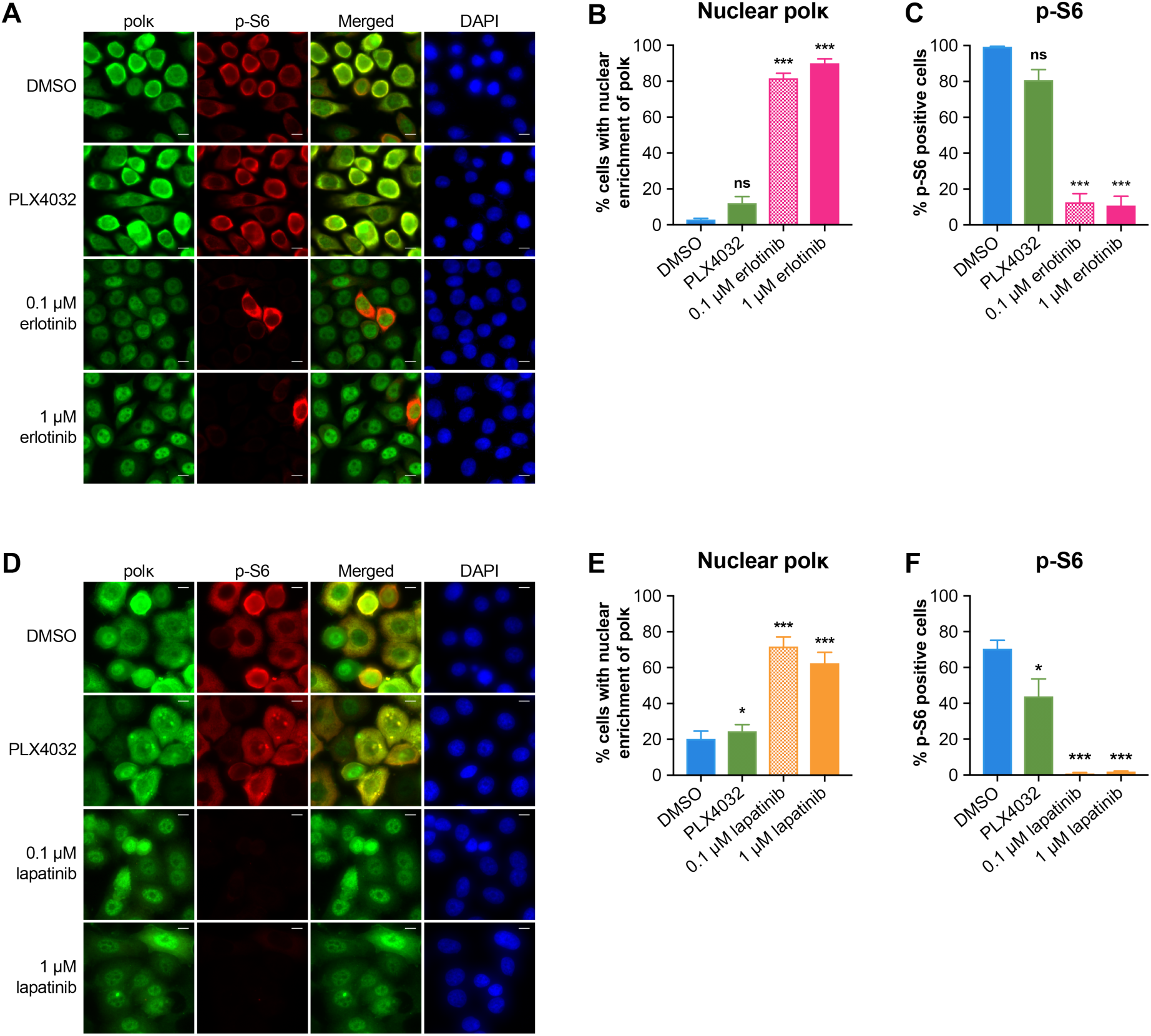
The effects on polκ and p-S6 are driver gene specific across tumor types. (**A**) Immunofluorescence staining of PC-9 cells treated with DMSO, 5 μΜ PLX4032, or erlotinib (0.1 μΜ or 1 μΜ) for 24 hours, scale bar 10 μm. (**B**) Quantification of the percentage of cells with nuclear enrichment of polκ, Mean ± S.E.M, at least 100 cells were counted for each sample across 4 fields of view, ns = non-significant, ***P < 0.001, paired two-tailed t-test. (**C**) Quantification of the percentage of cells with phosphorylated S6 (p-S6, S240/244), Mean ± S.E.M, at least 100 cells were counted for each sample across 4 fields of view, ns = nonsignificant, ***P < 0.001, paired two-tailed t-test. (**D**) Immunofluorescence staining of SK-BR3 cells treated with DMSO, 5 μM PLX4032, or lapatinib (0.1 μM or 1 μM) for 24 hours, scale bar 10 μm. (E) Quantification of the percentage of cells with nuclear enrichment of polκ, Mean ± S.E.M, at least 200 cells were counted for each sample across 7 fields of view, *P < 0.05, ***P < 0.001, paired two-tailed t-test. (F) Quantification of the percentage of cells with phosphorylated S6 (p-S6, S240/244), Mean ± S.E.M, at least 200 cells were counted for each sample across 7 fields of view, *P < 0.05, ***P < 0.001, paired two-tailed t-test.

### Glucose starvation phenocopies the effects on subcellular localization of polκ

mTOR could elicit the observed effect on polκ through multiple downstream mechanisms, but because *E. coli* use the starvation response as part of DinB upregulation/activation, we wondered whether by inhibiting mTOR, we were mimicking a starvation response. To test this idea, we examined the effect of nutrient deprivation on polκ localization. We grew A375 melanoma cells in a variety of media conditions in which we selectively removed serum, glucose, or glutamine and measured polκ localization by immunofluorescence. Whereas we found that both glutamine and serum deprivation had small effects on nuclear polκ localization, glucose starvation resulted in a highly significant induction of nuclear polκ (Fig. S7, A and B). Because glucose is a major carbon source for rapidly growing cancer cells, this may be analogous to the starvation response induced in *E. coli* by deprivation of lactose, which is an important carbon source for bacteria.

### Exportin-1 plays a role in regulating the subcellular localization of polκ

This data led us to ask what mechanisms the cells may use to control cytoplasmic versus nuclear localization of polκ. A previous analysis of the protein structure of polκ demonstrated that the full-length protein contains a bipartite nuclear localization signal (NLS) towards the 3’ end of the gene (*30, 33*), and computational analysis (*58*) also indicated a likely nuclear export signal (NES). Deletion of the NLS region results in the protein completely localizing to the cytoplasm (*33*), which suggested to us that the localization results we saw above might be controlled by the nuclear import and/or export machinery. To test this, we utilized inhibitors of importin-β or exportin-1 (CRM1) and tested whether these affected the shift of polκ in response to BRAF inhibition in melanoma cells. We found that importazole (*59*), which inhibits importin-β, had little effect on polκ localization (Fig. S8, A and B). In contrast, co-treatment with leptomycin B (*60*), which inhibits exportin-1, accelerated the rate at which polκ localized to the nucleus: whereas 3 hours of PLX4032 typically only leads to ~30% induction of nuclear polκ, in the presence of leptomycin B, this increased to 60% (Fig. 4, A and B). In addition, we noted that leptomycin B alone (even in the absence of PLX4032) induced a small but significant increase in nuclear polκ, indicating that polκ normally cycles between the cytoplasm and nucleus. To further test the contribution of the nuclear export machinery, we performed washout experiments. A375 cells were treated with PLX4032 (to induce nuclear polκ) and then the drug was washed away (Fig. 4C). After washout, polκ relocates to the cytoplasm within 24 hours. However, if leptomycin B is added after PLX4032 washout, polκ remains present in the nucleus (Fig. 4, D and E). This data indicates that oncogenic signaling regulates the nuclear localization of polκ (at least in part) through the export machinery.

**Fig. 4.**
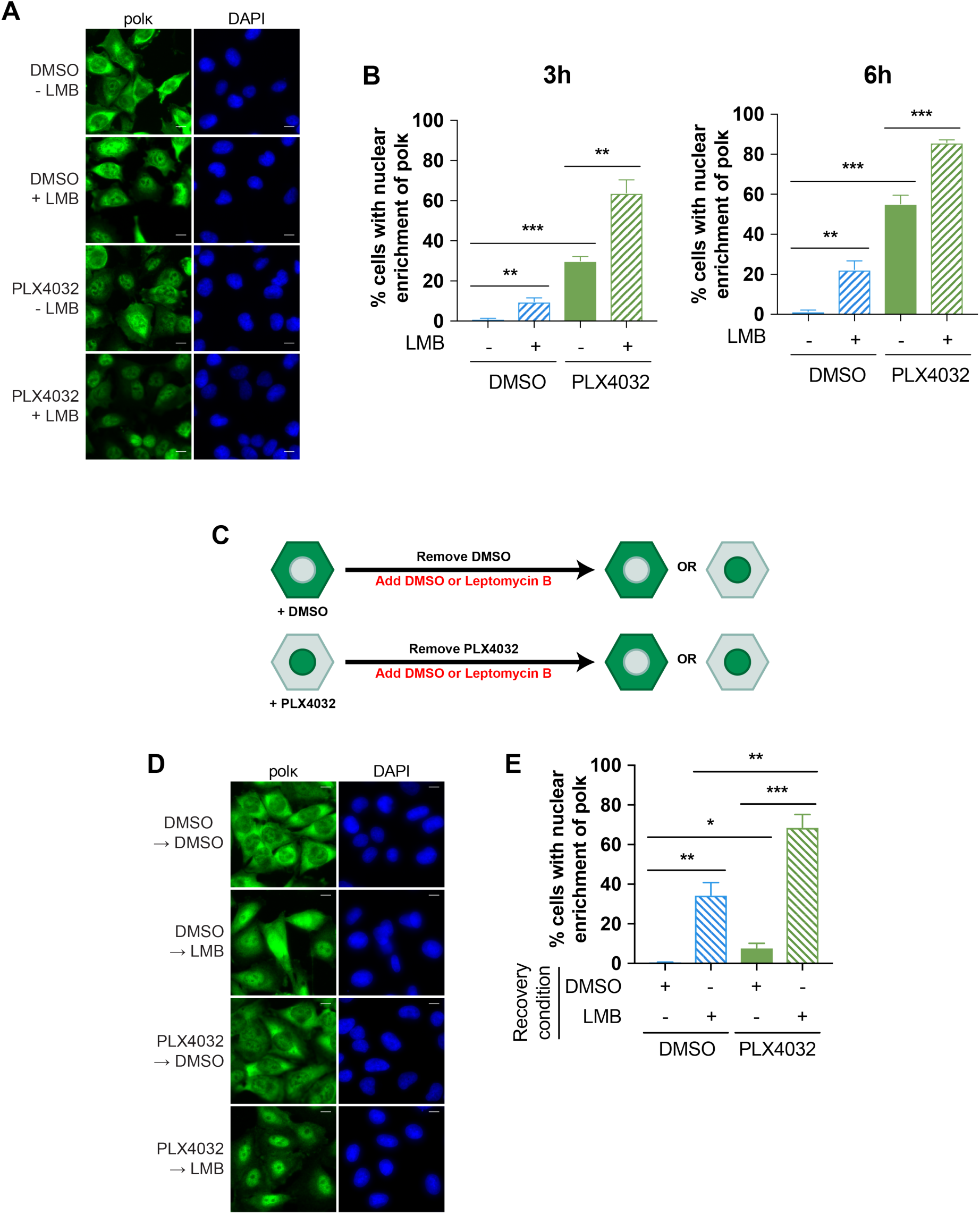
Exportin-1 plays a role in the subcellular localization of polκ. (**A**) Immunofluorescence staining of A375 cells treated with DMSO or 5 PLX4032 ± 20 leptomycin B (LMB) for 6 hours, scale bar 10 μm. (**B**) Quantification of the percentage of cells with nuclear enrichment of polκ after 3 or 6 hours, Mean ± S.E.M, at least 100 cells were counted for each sample across 4-8 fields of view, **P < 0.01, ***P < 0.001, paired two-tailed t-test. (**C**) Schema detailing the experiment design for the leptomycin B (LMB) recovery assay (D,E). (**D**) Immunofluorescence staining of A375 cells treated with DMSO or 5 μM PLX4032 for 24 hours and then DMSO or 10 μM LMB for 24 hours, scale bar 10 μm. (**E**) Quantification of the percentage of cells with nuclear enrichment of polκ, Mean ± S.E.M, at least 100 cells were counted for each sample across 7 fields of view, *P < 0.05, **P < 0.01, ***P < 0.001, paired twotailed t-test.

### polκ overexpression can lead to increased drug resistance

These data suggested a model in which rapidly growing cancer cells generally keep polκ at relatively low levels and sequestered in the cytoplasm due to active nuclear export. This is likely important to prevent excess mutagenesis in the absence of a need for translesion synthesis. Consistent with this idea, it has been previously shown that forced overexpression of polκ leads to sustained nuclear expression, and that this overexpression can be highly mutagenic on its own, i.e. even in the absence of overt DNA damage (*32*). We hypothesized that subverting this normal shuttling and forcing polκ to remain nuclear could contribute to drug resistance due to this capacity for inducing new mutations. To test this, we generated a doxycycline-inducible polκ overexpression construct for use in A375 melanoma cells (Fig. 5A). We then isolated multiple single cell clones from this population and expanded them in culture in the presence or absence of doxycyline for 3 months. As expected, these cells showed strong induction with increased nuclear localization of polκ after addition of doxycycline (Fig. S1, B and C). We then tested both populations for sensitivity to either the BRAF^V600E^inhibitor PLX4032 or the CDK4/6 inhibitor PD0332991 (Fig. 5A). For both clones, the population that had experienced long-term polκ overexpression showed a modest increase in growth in PLX4032 across multiple different doses compared to their respective negative control cells (Fig. 5B), consistent with increased resistance to the drug. In contrast, there was no significant difference between the two populations with regards to their responses to PD0332991 (Fig. 5C). This is consistent with the concept that prolonged nuclear expression of polκ is associated with resistance to certain clinically relevant cancer therapeutics, such as BRAF inhibitors.

**Fig. 5.**
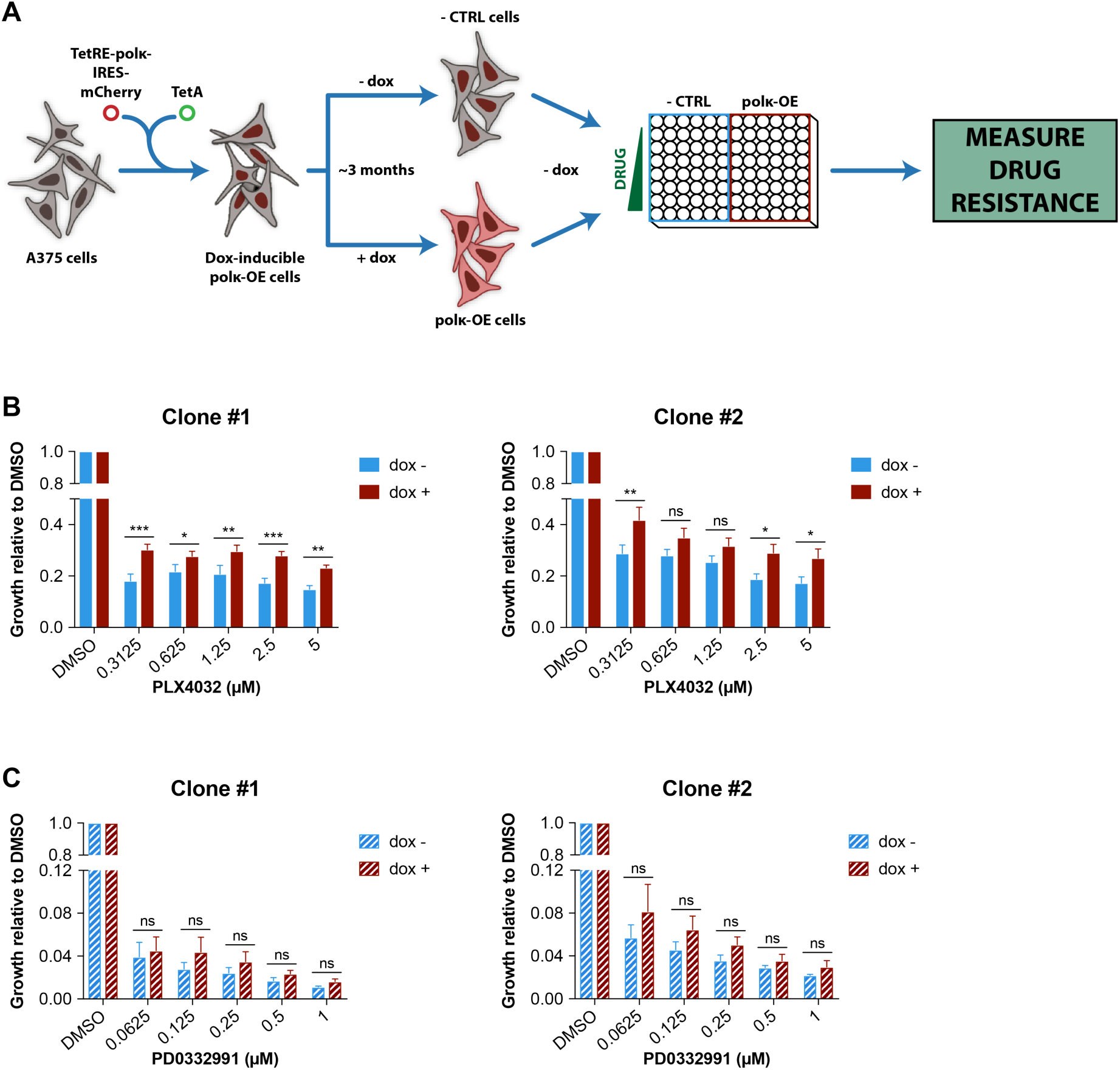
polκ overexpression can lead to increased drug resistance. (**A**) Schema detailing how single cell clones of A375 cells containing the doxycyline (dox)-inducible polκ overexpression construct were created and how the dox- and dox+ populations were generated and then used to measure drug resistance. (**B**) Drug resistance of the dox- and dox+ populations of two different clones of dox-inducible polκ overexpression cells to PLX4032 was determined by the CyQuant Direct assay. For each population, the relative cell viability at each dose, compared with the 0 μM dose control (DMSO), was calculated. Mean ± S.E.M, n=4-5 experiments, ns = non-significant, *P < 0.05, **P < 0.01, ***P < 0.001, Fisher's Least Significant Difference (LSD) test. A comparison between the two populations in their response to PLX4032 by two-way ANOVA gave a p-value of 0.0130 for Clone #1 and of 0.0294 for Clone #2. (C) Drug resistance of the dox- and dox+ populations of two different clones of dox-inducible polκ overexpression cells to PD0332991 was determined by the CyQuant Direct assay. For each population, the relative cell viability at each dose, compared with the 0 μM dose control (DMSO), was calculated. Mean ± S.E.M, n=3-4 experiments, ns = non-significant, Fisher's Least Significant Difference (LSD) test. A comparison between the two populations in their response to PD0332991 by two-way ANOVA gave a p-value of p = 0.3785 for Clone #1 and of p = 0.2278 for Clone #2.

## Discussion

There have been multiple reports of how alterations to DNA replication and repair processes can contribute to tumor initiation and progression. For example, defects in mismatch repair proteins such as MLH1 (*61, 62*), MSH2 (*63*), MSH6 (*64*), or PMS2 (*65*) are associated with Lynch syndrome, which predisposes patients to a wide variety of tumors including colon and gynecologic cancers (*66, 67*). More recently, large scale genome sequencing efforts have identified point mutations in the replicative polymerases polδ or polε, which inactivate the proofreading capacity of these proteins and induce a large number of mutations in the tumors that emerge (*1–3*).

Several studies have now identified overexpression of polκ in human tumors such as lung cancer or glioblastoma^9,10^. Because polκ lacks a proofreading domain, its overexpression can be associated with increased mutation rates as well as drug resistance (*11, 12*). However, despite the potential importance of polκ in promoting tumor progression, little is known about the mechanisms by which it is regulated. In this study, we identified oncogenic signaling and nutrient status as one potential regulator of polκ expression and localization in cancer.

polκ belongs to a family of related Y-family DNA polymerases which all function in translesion synthesis, a major DNA damage tolerance pathway (*7, 8*). In normal physiology, these polymerases play an important role in bypassing stalled replication forks that are induced by DNA damaging agents. Because they all lack proofreading domains, they have a propensity to introduce errors during replication, although depending on the specific lesion, they can also act in an error-free manner. For this reason, these Y-family polymerases can be a double-edged sword since they allow for replication past damaged regions of DNA and cell survival but may do so at a cost of new mutations (*68*). *in vitro* studies have demonstrated that these polymerases can act with extraordinarily low fidelity, with error rates ranging from 10^−1^ to 10^−4^, compared to ~10^−6^ for the normal replicative polymerases polδ or polε (*16, 69, 70*). For this reason, it is important that cells regulate the expression and localization of these polymerases such that they only act when the cell is under genomic stress.

Although DNA damage is likely the major inducer of polκ, our data would suggest that mammalian cells harbor the capacity to induce the expression of polκ under other forms of cell stress, namely loss of oncogenic signaling and/or nutrient starvation. This may be analogous to *E. coli*, whereby cells under starvation stress can upregulate DinB/pol IV (the bacterial orthologue of polκ) (*19, 20*). However, in *E. coli*, in addition to activating DinB, the RpoS and SOS pathways have other effects on the cells, including suppression of mismatch repair (*29*), and all of these processes are necessary to induce new mutations (*22*). Additionally, previous work has demonstrated that cancer cells harboring mutations in the proofreading domain of polε also experience loss of mismatch repair enzymes (*71*), and more recently it has been shown that suppression of mismatch repair is necessary for these polε mutants to dramatically increase their mutation rates (*72*). Surprisingly, in mammalian cells, overexpression of polκ alone has been shown to be highly mutagenic (*73*), although the status of mismatch repair was not directly assessed. Although in our studies we did not directly measure mutation rate after prolonged overexpression of polκ, we do see increased resistance to BRAF inhibitors, which is sometimes linked to the emergence of new clones with distinct mutational profiles (*74–80*). These mutations are often, although not always, found to be pre-existing upon deep sequencing (*81*). However, it is important to note that we cannot exclude other, non-mutagenic functions of polκ that could account for the increased drug resistance, the elucidation of which will require future evaluation.

One unexpected finding in our study was the observation that polκ can exist in a cytoplasmic form. Numerous prior studies have shown that polκ is primarily a nuclear protein, but these relied upon overexpression of an EGFP-polκ fusion protein (*32, 33*). In our hands, overexpression of polκ also strongly upregulated nuclear expression, which we believe likely reflects saturation of the export machinery. This would be consistent with our data using leptomycin B, which suggested that one important mechanism of regulation for polκ is nuclear-cytoplasmic shuttling. A few other studies have also observed scant cytoplasmic polκ in HeLa cells, including data from the Human Protein Atlas (*42*), which used antibodies raised against peptide antigens. One important difference in our studies is that we used a monoclonal antibody raised against the full-length protein. Examination of polκ transcript variants in Ensembl (*82*) reveals that humans transcribe at least 2 versions of polκ (polκ-201 and polκ-216), such that antibodies raised against short peptides versus full-length protein may recognize different transcripts with different localizations. The exact reasons for this discrepancy will await further studies of polκ structure in regards to post-translational modifications of nuclear localization/export signals or variations in transcript abundance in different cell types. Interestingly, another Y-family polymerase polη has also been reported to have nuclear localization when overexpressed as an EGFP fusion protein (*83*), yet data from the Human Protein Atlas (*42*) shows a predominantly cytoplasmic localization with nuclear enrichment in some cells, similar to what we see with polκ.

Our data indicates that mTOR and nutrient sensing regulate the localization of polκ, and this pathway more broadly is known to affect nuclear-cytoplasmic shuttling of multiple proteins (*84*– *91*). For example, PI3K/Akt/mTOR signaling has been shown to prevent nuclear accumulation of glycogen synthase kinase 3β (GSK3β) such that inhibition of this pathway leads to a robust increase in nuclear GSK3β (*89, 90*). Furthermore, mTORC1-dependent phosphorylation of the transcription factor TFEB promotes association of TFEB with members of the 14-3-3 family of proteins, thereby forcing its retention in the cytosol(*88*). However, the network connecting the mTOR pathway to the DNA damage response continues to be elucidated (*92–95*). A recent study (*95*) showed that mTOR/S6K signaling leads to phosphorylation and subsequent degradation of RNF168, and hyperactivation of mTOR via LKB1 leads to decreased levels of RNF168 and increases DNA damage. One possible explanation for our observed results is that mTOR (or one of its downstream targets) could directly phosphorylate polκ and thereby affect its localization as has been shown for other proteins that undergo nuclear-cytoplasmic shuttling (*96–99*). An important area for future exploration is understanding the ways in which mTOR may interact with the maintenance of genome stability.

One implication of our study is that polκ may play a role in resistance to targeted therapies. A previous study (*12*) showed that polκ could mediate resistance to the DNA damaging agent temozolomide, but no studies have directly linked polκ to non-mutagenic therapies such as BRAF^V600E^inhibitors. Both genetic and non-genetic mechanisms of BRAF inhibitor resistance have been described (*74–81, 100, 101*), and many of the mutations that lead to drug resistance are pre-existing in the population when examined by deep sequencing. However, it is also possible that certain forms of cell stress could lead to prolonged overexpression of polκ, resulting in new mutations in the population. Given the fact that the resistance phenotype we saw from such overexpression was modest, it is highly likely that there would need to be other defects in DNA repair for this to become a major mechanism of drug resistance. Such a mechanism may have similarities to the *E. coli* stress-induced mutagenesis pathway, in which overexpression of DinB/pol IV, in concert with other factors such as downregulation of mismatch repair, can lead to new drug resistance mutations (*19–23*). Given the availability of small molecule inhibitors of polκ (*102, 103*), it will be of future interest to determine whether the administration of such inhibitors could forestall the development of resistance to targeted therapies.

## Materials and Methods

### Cell culture

Cell culture was performed at 37°C in a humidified atmosphere containing 5% CO_2_. A375 and SK-MEL28 cells were obtained from ATCC. PC-9 cells were a gift from C. Rudin (MSKCC, NYC, USA). SK-BR3 cells were a gift from S. Chandarlapaty (MSKCC, NYC, USA). Cells were cultured in DMEM (A375 and SK-MEL28), RPMI1640 (PC-9), or DMEM/F12 (SK-BR3) supplemented with 2 mM glutamine, 100 IU/ml penicillin, 100 µg/ml streptomycin, and 10% heat-inactivated fetal bovine serum. Cell lines were regularly tested and verified to be mycoplasma negative by the MycoAlert™ Mycoplasma Detection Kit (Lonza).

### Cell treatments

Inhibitors were maintained until collection unless otherwise noted: PLX4032 (Selleck), CI-1040 (Selleck), U0126 (Sigma Aldrich), Ulixertinib (Selleck), SCH772984 (Selleck), erlotinib (Selleck), rapamycin (Selleck), PP242 (Abcam), LY294002 (Sigma Aldrich), bleomycin (Sigma Aldrich), importazole (Sigma Aldrich), and leptomycin B (Sigma Aldrich). Lapatinib and PD0332991 were gifts from S. Chandarlapaty. Complete media consisted of DMEM (without glucose or glutamine) supplemented with 25 mM glucose, 6 mM glutamine, and 10% heat-inactivated fetal bovine serum. Media – glucose consisted of DMEM (without glucose or glutamine) supplemented with 6 mM glutamine and 10% heat-inactivated fetal bovine serum. Media – glutamine consisted of DMEM (without glucose or glutamine) supplemented with 25 mM glucose and 10% heat-inactivated fetal bovine serum. Media + 1% serum consisted of DMEM (without glucose or glutamine) supplemented with 25 mM glucose, 6 mM glutamine, and 1% heat-inactivated fetal bovine serum.

### Generation of inducible polκ overexpression cells

For inducible overexpression of polκ, the human *polκ* open reading frame was amplified from a constitutive polκ overexpression plasmid that was a gift from J.S. Hoffmann (Toulouse, France) and cloned using the In-Fusion Cloning kit (Clontech) into the pSIN-TREtight-MCS-IRES-mCherry-PGK-Hygro vector, which has a doxycycline-inducible promoter and adds an IRES-mCherry to the C-terminus of polκ, to generate pTRE-hpolk-IRES-mCherry. The pSIN-TREtight-MCS-IRES-mCherry-PGK-Hygro vector and its corresponding tet-activator (rtTA3) RIEP vector were gifts from S. Lowe (MSKCC, NYC, USA). The following primers were used for amplification:

TRE-hpolk F1 FW: CGGTACCCGGGGATCCCACCATGGATAGCACAAAG TRE-hpolk F1 RV: TTAGTCTTCGCGGCCGCTTACTTAAAAAATATATCAAGGG

Lentivirus was produced by transfection of HEK-293T cells with pTRE-hpolk-IRES-mCherry and the packaging plasmid psiAmpho (which was a gift from R. Levine, MSKCC, NYC, USA) at a 1:1 ratio. Transfection was performed using Fugene (Promega) reagent. The viral supernatant was collected 48 and 72 hours following transfection, filtered through a 0.45 µm filter (Thermo Fisher Scientific), and added to A375 cells with 8 µg/ml polybrene (Thermo Fisher Scientific). Infected cells were selected by treatment with 1 mg/ml hygromycin (Thermo Fisher Scientific). Lentivirus for RIEP was produced the same way and then used to infect A375 cells already containing pTRE-hpolk-IRES-mCherry. Doubly infected cells (pTRE-hpolk-IRES-mCherry/RIEP) were selected by treatment with 10 µg/ml puromycin (Thermo Fisher Scientific) and 1 mg/ml hygromycin. Single cell clones were then isolated, and their induction efficiency was tested via flow cytometry. Clones #1 and #2 showed the best induction after addition of 1 µg/ml doxycycline (Sigma Aldrich) compared to uninduced controls, so they were selected for future experiments.

### Inducible shRNA knockdown of polκ and p53

For inducible knockdown of polκ, two different shRNAs (*104*) targeting polκ were generated using PCR. The two polκ shRNAs (sh-polκ-1 and sh-polκ-2) were each cloned using restriction digestion and ligation into the LT3GEPIR vector (*105*), which allows doxycycline-inducible expression of a shRNA and GFP. The following primers were used for amplification:

miRE_XhoF: TACAATACTCGAGAAGGTATATTGCTGTTGACAGTGAGCG miRE_EcoRI: TTAGATGAATTCTAGCCCCTTGAAGTCCGAGGCAGTAGGCA

The following oligos were used as the templates for amplification:

sh-polκ-1: TGCTGTTGACAGTGAGCGCAAGAAGAGTTTCTTTGATAAATAGTGAAGCCACAGATGTATTTATCAAAGAAACTCTTCTTATGCCTACTGCCTCGGA sh-polκ-2: TGCTGTTGACAGTGAGCGCAAGGATTTATGTAGTTGAATATAGTGAAGCCACAGATGTATATTCAACTACATAAATCCTTATGCCTACTGCCTCGGA

Plasmids containing either a control hairpin (sh-Ctrl) or 2 different hairpins targeting p53 (sh-p53-1 and sh-p53-2) were gifts from S. Lowe. For all hairpins, lentivirus was produced by transfection of HEK-293T cells with each hairpin plasmid with the packaging plasmids psPAX2 and pMD2G at a 4:3:1 ratio. Transfection was performed using Lipofectamine 2000 (Life Technologies) reagent. The viral supernatant was collected at 48 and 72 hours following transfection and frozen at -80°C. Virus was later thawed and added to A375 cells, and infected cells were selected by treatment with 1 µg/ml puromycin. Induction after treatment with 1 µg/ml doxycycline was verified using flow cytometry.

### Total RNA extraction, cDNA isolation and qRT-PCR analysis

Total RNA from treated cells was extracted using the Quick-RNA Mini-Prep kit (Zymo), and RNA concentration was determined using a Synergy H1 Hybrid Multi-Mode Microplate Reader (BioTek). Reverse transcription of total RNA was performed using SuperScript III First-Strand Synthesis SuperMix (Thermo Fisher Scientific) according to the manufacturer’s guidelines. qRT-PCR reactions were detected on a CFX384 Touch machine (Bio-Rad) using iQ SYBR Green Supermix (Bio-Rad). RNA expression levels were calculated using the comparative Ct method (2-^ΔΔCT^) normalized to β-actin. The primer pairs used were as follows:

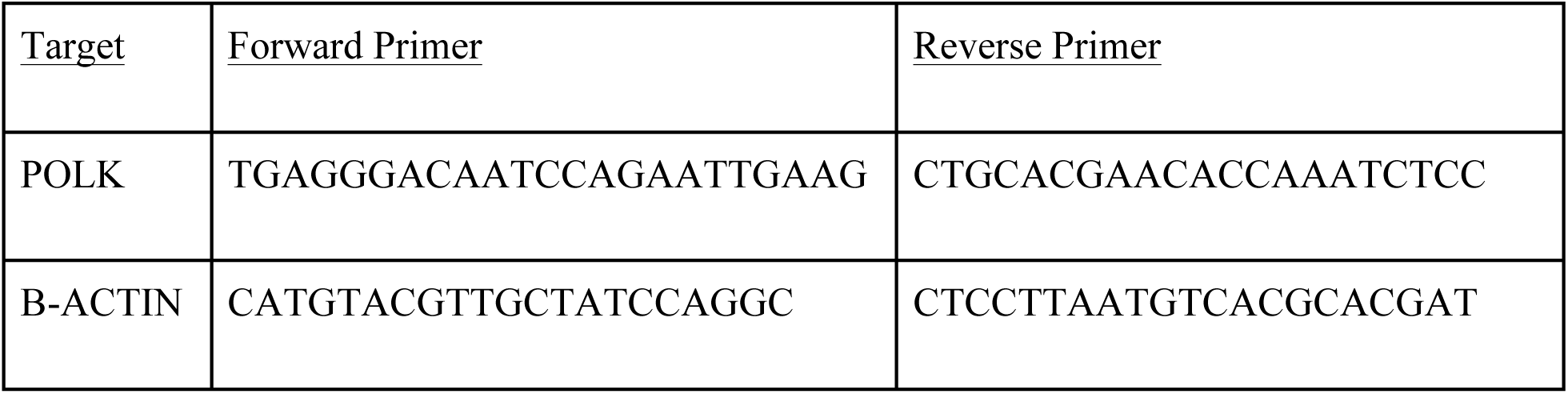

### Western blot analysis

For whole cell lysates, cells were washed 1x with phosphate buffered saline (PBS, Invitrogen) and then lysed in RIPA buffer (EMD Millipore) containing 1x HALT Combined Protease and Phosphatase Inhibitor Cocktail (Thermo Fisher Scientific) for 20 minutes at 4°C. Cell lysates were clarified by centrifugation at 10,000 RPM for 10 min at 4°C. For cytoplasmic or nuclear fractions, the Subcellular Protein Fractionation Kit for Cultured Cells (Thermo Fisher Scientific) was used according to the manufacturer’s protocol. Protein concentration was measured with Bradford reagent (Sigma Aldrich), and samples were resolved by 4–15% or 12% SDS-PAGE gels (Bio-Rad) for polκ or p53, respectively. Proteins were transferred onto nitrocellulose membranes and subjected to standard immunoblotting. ECL Prime (Amersham) was used as the developing reagent.

*Western blot antibodies:* polκ (ab57070, Abcam, 1:10,000), p53 (sc-126, Santa Cruz Biotechnology, 1:10,000), mcherry (ab167453, Abcam, 1:1000), lamin-B1 (ab133741, Abcam, 1:10,000), β-Actin (A5441, Sigma Aldrich, 1:20,000), anti-mouse (ab97046, Abcam, 1:10,000), anti-rabbit (ab97051, Abcam, 1:10,000)

### Immunofluorescence

Cells were cultured on Millicell EZ SLIDE 4 or 8-well glass slides (EMD Millipore). Cells were fixed with 4% paraformaldehyde (Santa Cruz Biotechnology) at 37°C for 15 minutes, washed with PBS, and then blocked with 5% goat serum (Thermo Fisher Scientific) and 0.2% Triton X-100 (Thermo Fisher Scientific) in PBS for 1 h. The cells were incubated with primary antibody in antibody dilution buffer (PBS with 1% bovine serum albumin (Sigma Aldrich) and 0.2% Triton X-100) at 4°C overnight, then washed 3 times with PBS, and incubated with Alexa-conjugated secondary antibody for 2 h at RT. After washing 3 more times with PBS, cells were stained with 0.1 µg/ml DAPI (Thermo Fisher Scientific) and then mounted with Dako fluorescence mounting media (Agilent) and imaged on a Zeiss Axio Imager A2.

*Immunofluorescence antibodies:* polκ (ab57070, Abcam, 1:500), p-S6 (5364, Cell Signaling, 1:1000), γH2AX (2577, Cell Signaling, 1:200), 53BP1 (ab175933, Abcam, 1:200), p-Chk1 (2348, Cell Signaling, 1:100), p-Chk2 (2661, Cell signaling, 1:500), anti-mouse Alexa-488 (4408, Cell signaling, 1:1000), anti-rabbit Alexa-488 (4412, Cell Signaling, 1:1000), anti-rabbit Alexa-594 (8889, Cell Signaling, 1:1000)

### Cell cycle analysis

Cells were treated with the indicated inhibitors for 24h and fixed in 70% ethanol (Thermo Fisher Scientific) at 4°C for at least 4 hours. Later, the cells were washed with PBS and stained with 20 µg of propidium iodide (Thermo Fisher Scientific), 200 µg of RNase A (Thermo Fisher Scientific), and 0.1% Triton X-100 in PBS for at least 30 minutes. The labeled cells were analyzed using a Fortessa flow cytometer, and the Dean-Jett Fox model in FlowJo was used to determine which cells were in G1, S, and G2/M.

### Drug resistance assays

Two different clones of doxycycline-inducible polκ overexpression cells (Clone #1 and Clone #2) were generated. We then divided each population equally and treated with or without 1 µg/ml doxycycline for ~3 months to generate polκ-overexpressing cells and their corresponding negative control population. All populations were then switched to media without doxycycline, plated in 96 well plates, and exposed to various doses of PLX4032 or PD0332991. After 4 (PLX4032) or 7 days (PD0332991), cell number was determined using the CyQUANT Direct Cell Proliferation Assay kit (Thermo Fisher Scientific). The technical replicates for each dose were averaged and then normalized against the 0 µM dose condition to determine the growth relative to DMSO.

### Statistical tests

The following statistical tests were used: paired 2-tailed t-test (all q-RT-PCR, Western blot analysis, and immunofluorescence), chi-square tests (cell cycle analysis), Pearson correlation coefficient (comparison of rates of p-S6 loss vs. nuclear polκ enrichment), and two-way ANOVA and Fisher’s least significant difference (LSD) test (drug resistance assays).

## Supplementary Materials

Fig. S1. Validation of the polκ antibody.

Fig. S2. MAPK inhibition in melanoma cell lines increases mRNA levels of Y-family polymerases and changes polκ’s subcellular localization.

Fig. S3. DNA damage is not associated with the subcellular shift of polκ.

Fig. S4. Cell cycle inhibition is not responsible for the shift in polκ’s subcellular localization.

Fig. S5. MAPK inhibition decreases phospho-S6.

Fig. S6. The effects on polκ mRNA levels are driver gene specific across tumor types.

Fig. S7. Glucose starvation also modulates polκ subcellular localization.

Fig. S8. Inhibiting Importin-β does not prevent the nuclear shift of polκ.

## Acknowledgements

We thank S. Chandarlapaty (MSKCC, NYC) for providing lapatinib and PD0332991; J.S. Hoffmann (Toulouse, France) for the constitutive polκ overexpression plasmid; S. Lowe (MSKCC, NYC) for the pSIN-TREtight-MCS-IRES-mCherry-PGK-Hygro vector, its corresponding tet-activator (rtTA3) RIEP vector, and plasmids containing either a control hairpin (sh-Ctrl) or 2 different hairpins targeting p53 (sh-p53-1 and sh-p53-2); and R. Levine (MSKCC, NYC) for the packaging plasmid psiAmpho. We also thank Wenjing Wu for help with the figures.

**Funding:** This work was supported by the Ruth L. Kirschstein Predoctoral Individual National Research Service Award (5F31CA200341-03) (K.T.), the NIH Director’s New Innovator Award (DP2CA186572), Mentored Clinical Scientist Research Career Development Award (K08AR055368), the Melanoma Research Alliance, The Pershing Square Sohn Foundation, The Alan and Sandra Gerry Metastasis Research Initiative at the Memorial Sloan Kettering Cancer Center, and The Harry J. Lloyd Foundation and Consano (all to R.M.W.).

**Author contributions:** K.T. and R.M.W. designed the study and wrote the manuscript. K.T. performed all experiments except as follows: K.M. performed the experiments on SK-MEL28 cells, A.C. and E.M.L. helped with some qRT-PCR and immunofluorescence experiments.

**Competing interests:** The authors declare that they have no competing interests.

